# fMRI evidence of equivalent neural suppression by repetition and prior knowledge

**DOI:** 10.1101/056069

**Authors:** J. Poppenk, M. Moscovitch, A.R. McIntosh

**Affiliations:** Department of Psychology, Queen′s University, 62 Arch St., Kingston, ON, Canada K7L 3N6; Department of Psychology, University of Toronto, 100 St. George St., Toronto, ON, Canada M5S 3G3; Rotman Research Institute, Baycrest, 3560 Bathurst St., Toronto, ON, Canada M6A 2E1

## Abstract

Stimulus repetition speeds behavioral responding (behavioral priming) and is accompanied by suppressed neural responses (repetition suppression; RS) that have been observed up to three days after initial exposure. While some proposals have suggested the two phenomena are linked, behavioral priming has been observed many years after initial exposure, whereas RS is widely considered a transitory phenomenon. This raises the question: what is the true upper limit of RS persistence? To answer this question, we scanned healthy, English-native adults with fMRI as they viewed novel (Asian) proverbs, recently repeated (Asian) proverbs, and previously known (English) proverbs that were matched on various dimensions. We then estimated RS by comparing repeated or previously known proverbs against novel ones. Multivariate analyses linked previously known and repeated proverbs with statistically indistinguishable RS in a broad visual-linguistic network. In each *suppressed* region, prior knowledge and repetition also induced a common shift in functional connectivity, further underscoring the similarity of the RS phenomenon induced by these conditions. By contrast, *activated* regions readily distinguished prior knowledge and repetition conditions in a manner consistent with engagement of semantic and episodic memory systems, respectively. Our results illustrate that regardless of whether RS is understood in terms of its magnitude, spatial extent or functional connectivity profile, typical RS effects can be elicited even under conditions where recently triggered biological processes or episodic memory are unlikely to play a prominent role. These results provide important new evidence that RS (of the kind observed after an interval of at least several minutes) reflects the facilitation of perceptual and comprehension processes by any type of information retrieved from long-term memory.

---

In some brain regions, responses to stimuli are suppressed after the first exposure (i.e., *RS*; a.k.a. *neural priming, neural adaptation, or novelty activation)*. This phenomenon occurs reliably in perceptual or semantic processing regions in human fMRI studies (Schacter, Wig, & Stevens, 2007) and may mediate more rapid and accurate responding to repeated stimuli (behavioural priming; Turk-Browne, Yi, & Chun, 2006). Behavioral priming, however, can persist for many years (Kolers, 1976; Mitchell, 2006; Sloman, Hayman, Ohta, Law, & Tulving, 1988), whereas RS is widely regarded as a transitory phenomenon. While studies report RS effect size does decline rapidly as the interval between exposure and repetition extends from seconds to minutes (Henson, Rylands, Ross, Vuilleumeir, & Rugg, 2004; Henson, Shallice, & Dolan, 2000; Wagner, Maril, & Schacter, 2000), RS can persist, nonetheless, for several days (Henson, 2003; van Turennout, Ellmore, & Martin, 2000), apparently without further decay (van Turennout, Bielamowicz, & Martin, 2003). These findings suggest that longer lasting RS, in contrast to the rapidly decaying one, reflects a relatively enduring change to cortical processing.

In the current study, we evaluated a remote upper bound for the durability of the type of neural suppression associated with repetition over intervals of more than several minutes. Our novel approach was to ask: can such neural suppression also be evoked by long-dormant stimuli which presumably had been repeated long ago? If neural suppression, of which RS is one type, reflects permanent changes in the brain, then presentation of stimuli encountered many years ago should elicit neural suppression substantially equivalent to that evoked by recently repeated stimuli (where *substantially equivalent* is defined broadly as having high overlap in features, such as in response magnitude, spatial distribution, and connectivity). In contrast, if neural suppression is transient, there should be little resemblance in the neural signature of repetition suppression induced by recently and remotely presented stimuli. We scanned native English-speaking participants using fMRI while presenting novel Asian proverbs, recently repeated Asian proverbs, and English proverbs previously known to participants (based on past repetitions through non-laboratory experience; Table 1).

**Table 1.**
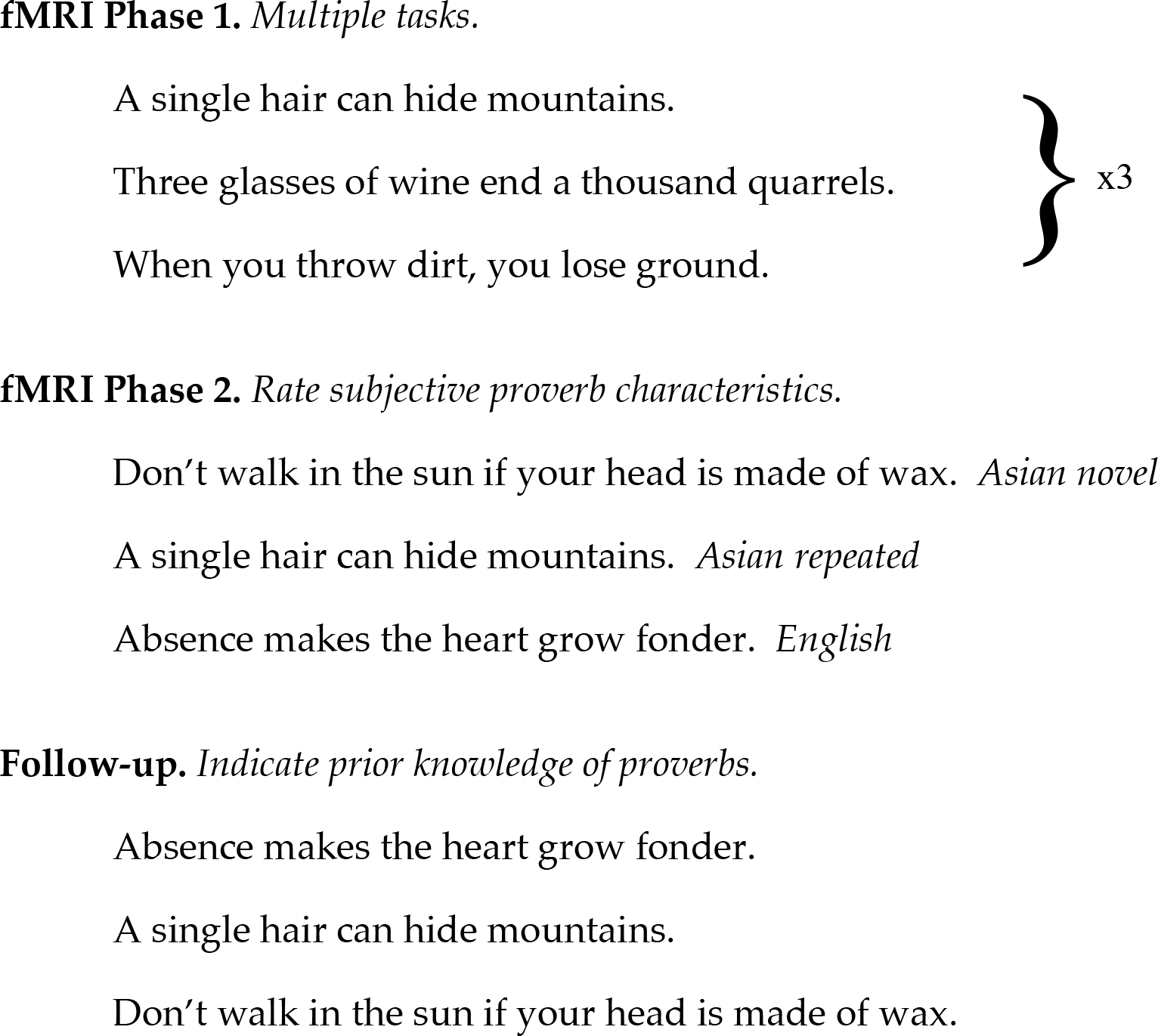
*Schematic of experimental protocol*. In Phase 1, participants viewed the Asian repeated list of proverbs three times. In Phase 2, they rated the repeated proverbs, a novel list of Asian proverbs, and a list of English proverbs in terms of subjective characteristics. In a follow-up session, participants confirmed which proverbs they knew prior to the experiment.

As RS has been reported in sensory and language cortices in response to repeated verbal material (Dehaene-Lambertz et al., 2006; Hasson, Nusbaum, & Small, 2006), we anticipated similar effects in these regions for recently repeated Asian proverbs. Of primary theoretical interest, however, was how RS for repeated Asian proverbs would compare to neural suppression induced by common English proverbs whose repetition occurred long before the participants entered the experiment. In generating specific hypotheses, it is important to consider not only the possible trajectory of decay, but also consolidation of longterm memory of our recent events and experiences (episodic memory) into our corpus of world knowledge (semantic memory), a phenomenon that most theoretical accounts posit to occur over longer time scales (Eichenbaum, Yonelinas, & Ranganath, 2007; Moscovitch et al., 2005; Squire & Zola, 1998). Because Asian proverbs were introduced about 30 minutes before they were repeated within the same experimental session, by most accounts, it is unlikely they had sufficient time to enter into semantic memory. By contrast, at any given time, it is unlikely English speakers will have encountered a particular English proverb within the past few months or years (see *Materials*); yet, most are familiar from a lifetime of exposures. As episodic memory is likely to decay over such intervals, presentation of English proverbs to English native-speakers is more likely to trigger implicit retrieval from semantic than episodic memory. Therefore, English proverbs and recently repeated Asian proverbs likely differed in that they were more likely to trigger implicit retrieval from semantic and episodic memory, respectively.

Turning to our hypotheses, we anticipated two possible outcomes concerning RS. To the extent that RS effects primarily reflect the availability of information from memory to guide perception and comprehension, RS might be the equivalent whether information comes from episodic or semantic memory. Accordingly, equivalent RS might be induced by repeated Asian proverbs and previously known English proverbs. By contrast, if RS effects primarily track the gradual decay of a representation triggered by recent exposure, RS might be present among recently repeated Asian proverbs, but largely absent when comparing novel proverbs against previously known English ones. In both cases, we expected that any RS effects would be localized principally within visual and linguistic regions.

Whether or not episodic and semantic memory are implicated in RS, both memory systems are thought to operate automatically, with retrieval events triggered independently of need or intention by the presentation of a pertinent cue (Moscovitch, 1992; Moscovitch, Cabeza, Winocur and Nadel, 2016). Consequently, their differing involvement in response to the two proverb types may reasonably be expected to lead to different amounts of retrieval from the
two memory systems. As a large number of experiments have shown, episodic and semantic memory rely on different neural systems from one another (Eichenbaum et al., 2007; Moscovitch et al., 2005; Squire & Zola, 1998), with episodic memory retrieval tasks tending to activate brain areas such as the hippocampus, precuneus, and lateral prefrontal cortex, and semantic memory retrieval tasks activating brain areas such as lateral temporal cortex and the temporal pole. Therefore, we predicted that the brain areas *activated* by English and repeated Asian proverbs (i.e., repetition enhancement; RE) are likely to be quite different, and associated with the sets of regions linked to semantic and episodic memory, respectively. We tested these ideas using both whole-brain multivariate analyses, and voxel-wise approaches.

As a means of evaluating RS for repeated materials and for prior knowledge materials, we separately compared fMRI responses to the repeated Asian and English proverbs against fMRI responses to novel Asian proverbs. It should be emphasized that this approach does not involve comparison of the neural signature of individual stimuli sampled at two points in time-this was impossible, as we had no reference measurement available for the English proverbs. Rather, our analysis rests upon the idea that the idiosyncratic features of individual stimuli are abstracted through random stimulus assignment across participants and the process of averaging across many items. By presenting items that are novel, repeated recently or previously known, we aimed to obtain a “typical” neural response profile for each class of item. With respect to both our hypotheses concerning RS and those concerning RE, it is worth noting that English and Asian proverbs could have subtle differences between them that contribute to activations and deactivations when they are directly contrasted; such differences, however, are unlikely to be mistaken for the specific outcomes predicted in our hypotheses. In particular, these hypotheses are: 1) RS will either appear similar for English and repeated Asian proverbs, if RS effects are permanent, or fail to appear at all for English ones, if RS effects require recent exposure; and that 2) RE will differ based on reliance of English and repeated Asian proverbs on semantic and episodic memory systems, respectively.

## Materials and Methods

### Participants

Eighteen right-handed volunteers from Toronto, Canada participated in the experiment. All were native English-speakers with normal or corrected-to-normal vision and hearing, (11 female; aged 21 to 34 years, mean age 26.1). All had less than one year of experience with East Asian languages and at least one native English-speaking parent or guardian. Participants were screened for neurological and psychiatric conditions and received financial remuneration. One participant was excluded for chance-level behavioural performance. The experimental protocol was approved by the Research Ethics Boards at the University of Toronto and Baycrest Hospital.

### Stimulus materials

We employed 160 Chinese and Japanese (Asian) proverbs. Each proverb consisted of a complete sentence of at least five words, and some were modified for smooth reading in English. Two 80-proverb lists were randomly generated for each participant: a ″repetition″ and ″novelty″ set. A third list of 80 common English proverbs was also compiled. It should be noted that proverbs have an extremely low frequency of occurrence in English discourse, with an approximate usage frequency of one in every 30 million to 200 million five-word phrases (within a large corpus of English books published 1990-2000; http://ngrams.googlelabs.com). Consequently, it is unlikely participants were exposed to any substantive number of English (or Asian) proverbs in the months before our experiment. The specific English and Asian proverbs we employed were selected based on participant knowledge in an earlier study (Poppenk, Kohler, & Moscovitch, 2010), and were matched for length (English *M* = 6.2 words, *SD* = 1.9 words; Asian *M* = 6.3 words, *SD* = 1.6 words). A fourth list of nonsense strings was prepared by randomly substituting the letters of each proverb.

### Procedure

The study consisted of two sessions: one involving two hours of fMRI scanning and one hour of computer tasks, and a second, completed within a day of the first, involving another hour of computer tasks. Prior to the main experiment, experimental tasks were practiced in a mock fMRI scanner using a six-item practice set. A schematic of this procedure is described in Table 1.

In the repetition phase (Phase 1), participants were scanned while viewing the “repetition” set of Asian proverbs three times in two tasks. In the first “deeper meaning” task, participants viewed each proverb and indicated with a button press when they derived a plausible meaning. Each of the 80 items from the repetition set was presented for 7.5 s with an inter-stimulus interval (ISI) of 4.0 to 10.5 s (average 5.9 s). Proverbs were displayed in a centered black 18-point bold Courier New font over a white background. Persistent text at the bottom of the screen in white text over a black banner (centered 12-point bold Courier New font) reminded participants of the response-key mapping and instructions. Each proverb was presented a second and third time in a “cultural origin” task designed to maximize familiarity with the proverbs through elaborative processing (Craik & Lockhart, 1972). Participants decided whether each proverb was South American or Japanese (all proverbs were Asian). During this task, proverbs were presented for 3.0 s with a 1.0 s ISI. Text formatting and a response-key mapping reminder were as in the first task. After all proverbs were presented, participants rated all proverbs a second time. The full task took 10.9 minutes. By the end of Phase 1, all 80 proverbs from the “repeated” Asian set had been presented three times. Participants then rested for 4.3 minutes while anatomical scans were completed.

In Phase 2, participants were scanned during four functional runs while subjectively rating qualities of novel Asian, English, or recently repeated Asian proverbs (Table 1 and Fig. 1). Participants rated each proverb in terms of its quality (“subjectively, did you find the proverb to be of high or low quality?”) or target audience (“Is this proverb better suited to a youth or an adult?”). We allocated different keys to each task so that use of the correct set of keys would be diagnostic of engagement in the correct rating task. A baseline task required participants to decide whether a nonsense string contained more x's or more o's (data from this task were not analyzed). Each of the four runs contained two superordinate blocks for each of the three tasks. Each of the six blocks was preceded by a cue designating the task to be performed (i.e., “target age”, “quality” or “x's and o's“). 4.5 s later, a key-mapping reminder was placed at the bottom of the screen and remained in place until the end of the block. To help distinguish the tasks and alert participants to changes in task, each task was allocated a color (green, navy or maroon) that was used for all text and the solid background color of the instruction bar. The color and keys assigned to each task were counterbalanced across participants. Within each target age and quality super-block, there was one sub-block containing five trials for each of the three proverb types (novel Asian, repeated Asian and English; for a total of fifteen trials per block). Likewise, each baseline block contained five trials.

**Figure 1.**
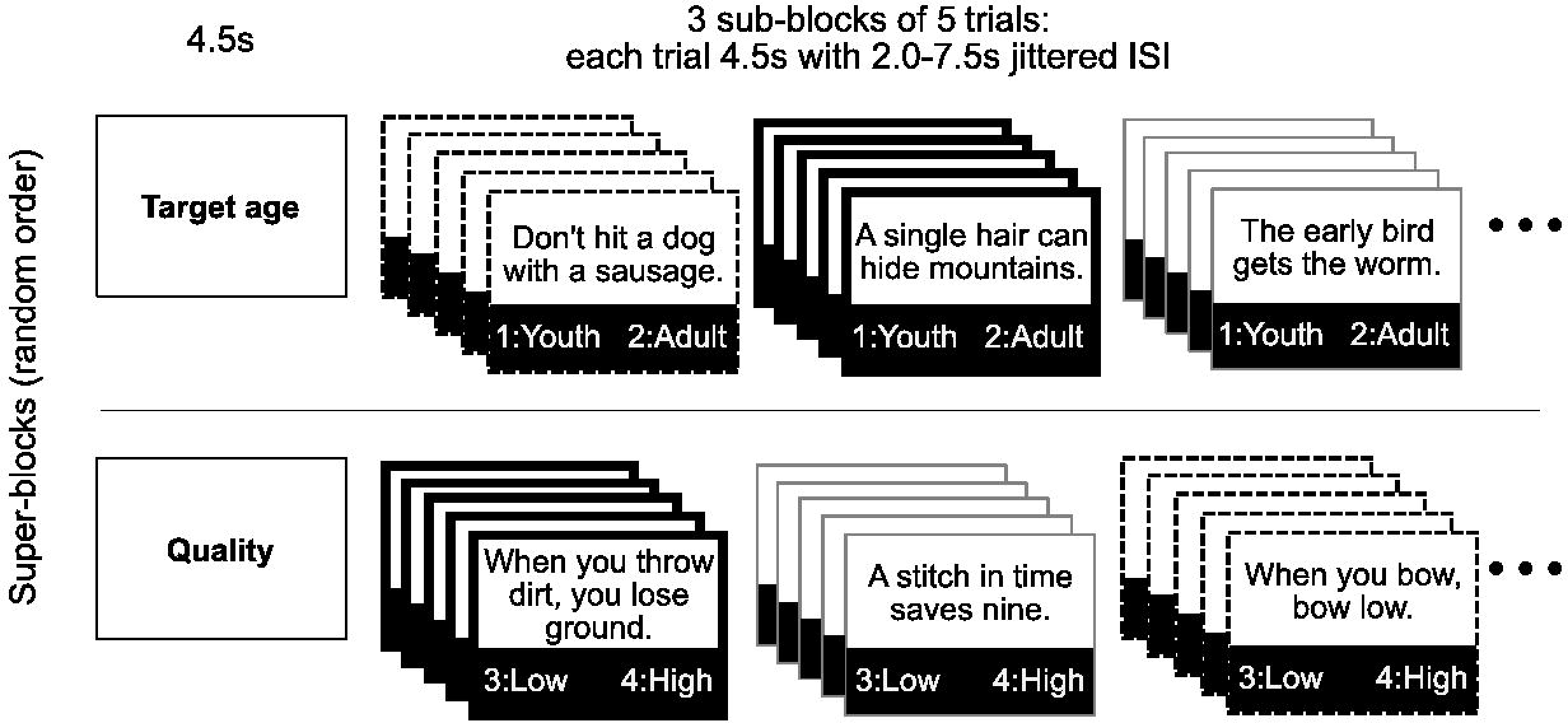
*Schematic of Phase 2 mixed design*. Participants completed incidental tasks on super-blocks consisting of three sub-blocks of five trials. Sub-blocks were novel (dashed line), known due to recent repetition (thick line), or known due to prior knowledge (thin line). The allocation of items to subblocks, the order of sub-blocks and the order of super-blocks were each randomized across participants.

In all blocks, trials began with the 4.5 s presentation of a proverb drawn randomly from the appropriate list (or a nonsense letter string in the case of baseline blocks), in centered text (18-point bold Courier New font over a white background). This presentation was followed by an inter-stimulus interval of 2.0 to 7.5 s (average 4.8 s). In total, each of the four runs in Phase 2 consumed 12.1 minutes in the scanner. Across all runs, 240 proverbs were presented-80 novel Asian, 80 repeated Asian and 80 English proverbs-with each type split evenly between the quality and target age tasks. Runs were separated by a 30-second break. The mean repetition interval from the third (final) exposure of proverbs repeated in Phase 1 to re-exposure in Phase 2 was 32.5 minutes (SD: 14.8, min: 5.6, max: 59.5).

For ten minutes following Phase 2, participants closed their eyes while fMRI resting state scans were acquired (no stimulation was administered). After scanning, participants completed a surprise memory test for all 240 Phase 2 items in a quiet room. Within 24 hours, participants scored their prior knowledge for the proverbs (Table 1), with all items from the three lists presented in random sequence. Participants were advised that all items had been presented, and that the task, therefore, was not a “memory test”, but a survey of their prior knowledge. They indicated with a button press whether each proverb was previously known or learned during the experiment (no response deadline was imposed). Each item was followed by a 1 s ISI. Stimuli and a response key-mapping reminder were presented as in Phase 1.

An overview of analyses performed on the behavioral and fMRI data we obtained is provided in Table 2.

**Table 2.**
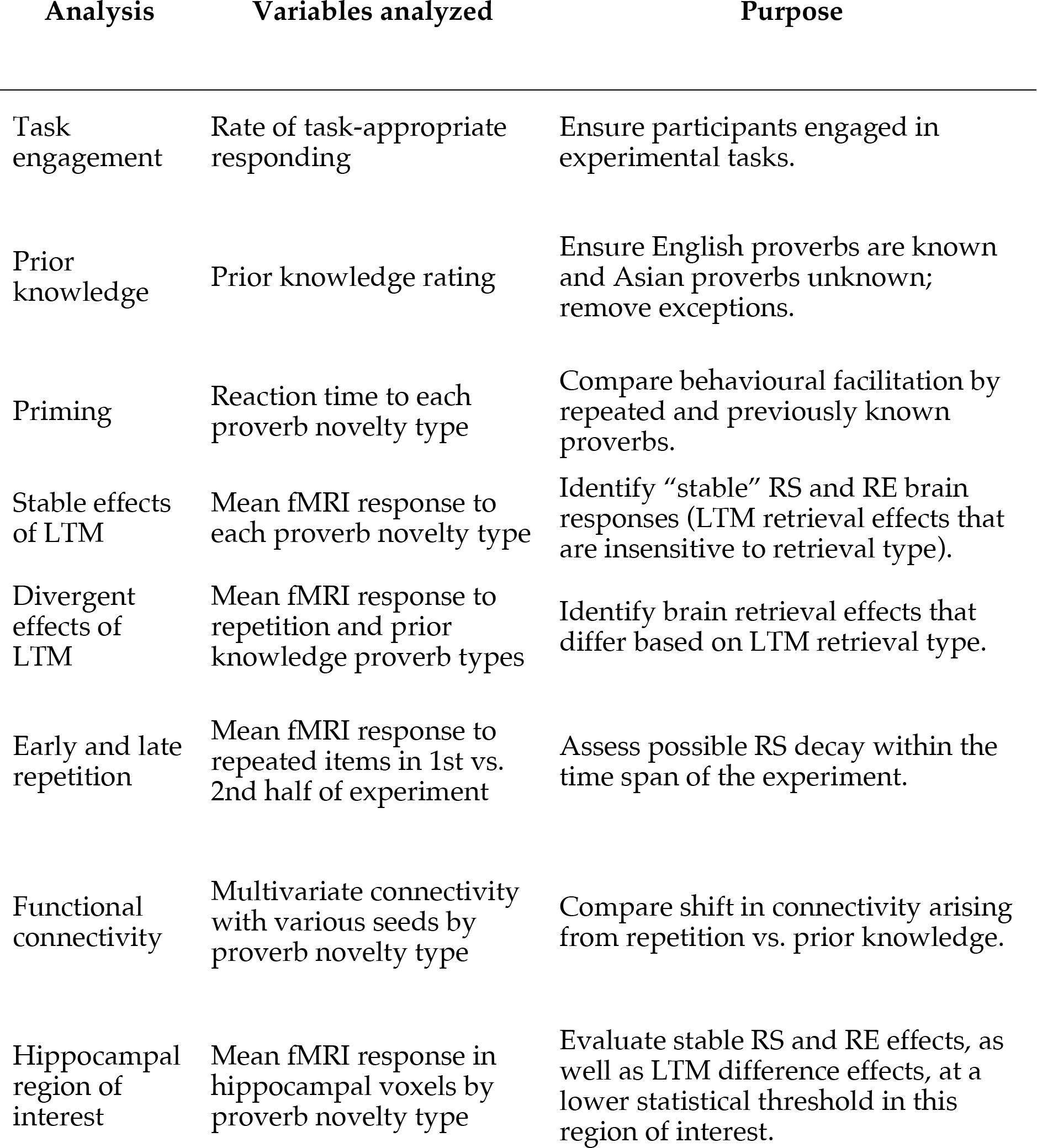
*Schematic of analyses performed*. Overview of analyses performed.

### Data acquisition and analysis

All imaging was performed on a 3 Tesla whole-body MRI system (Siemens, Erlangen, Germany) using a 32-channel array head coil. T2-weighted echo planar image acquisition was used for all functional scans (TE = 30 ms; TR = 1500 ms; flip angle = 60ΰ). Each functional run involved acquisition of ten discarded stabilization volumes and 475 task volumes. 24 contiguous 5mm-thick axial oblique slices captured the full neocortical brain volume of each participant, omitting the lower brain stem and inferior portion of the cerebellum (FOV = 200 × 200 mm; 64 × 64 matrix). A T1-weighted high-resolution MRI volume was also obtained using a 3D MPRAGE pulse sequence (160 slices; FOV = 256 × 256 mm; 192 × 256 matrix).

Initial preprocessing of the T2-weighted functional images was performed on a run-by-run basis using FSL (FMRIB Software Library version 4; Smith et al., 2004). Following correction for timing differences between slices in the same volume and motion correction within each image series, high-pass filtering was applied to exclude scanner drift and other low-frequency noise (sigma = 49.5 s). Each series was registered to the participant's high-resolution T1-weighted anatomical image and segmented to isolate the region containing the brain volume. Using the anatomical image as a reference, functional data were resampled into MNI space (Cocosco, Kollokian, Kwan, & Evans, 1997) using isotropic voxels (4 × 4 × 4 mm^3^). Each image series was then entered into a probabilistic independent component analysis, which was conducted on a run-by-run basis to identify and remove residual motion artifacts, high-frequency scanner noise and artifacts related to gradient timing errors. This step was performed using MELODIC (Beckmann & Smith, 2004). Two raters inspected components from one series of each participant. Their evaluations were used to train a component-sorting classification model, which evaluated remaining components, with manual spot checks made to verify sorting quality. Next, the data were despiked, and mean white matter intensity for each image volume was calculated across white matter voxels (identified using an MNI probability image) for residualization as a factor of no interest. Finally, images were z-scored over time and spatially smoothed using a 3D Gaussian kernel with a full-width at half maximum value of 6 mm.

Image series were parsed into event epochs of eight scans. These were combined to obtain a mean image for each condition and participant. The only difference between this approach and the procedure we used for our functional connectivity analyses was that in the latter case, for each condition and participant, we created a correlation image instead of a mean image. We did so by obtaining, within every epoch, the time series associated with a seed voxel in that condition and participant. Then, also within each epoch, we independently regressed the seed time series with the time series of every other voxel in the brain. This yielded a correlation coefficient image containing a different coefficient in every voxel for every epoch, reflecting the coactivation of that voxel with the seed. We then averaged the correlation images across epochs to create a mean correlation image for each condition and participant, which was then entered into further analysis.

All fMRI analyses were conducted using non-parametric resampling statistics. Pairwise hypothesized contrasts were performed using non-rotated partial least squares in PLSGUI (McIntosh & Lobaugh, 2004). Non-rotated PLS is a hypothesis-driven analysis suitable for testing whether hypothesized contrasts are associated with stable large-scale activity patterns, and for evaluating which brain voxels are reliably associated with the contrasts. For each analysis, we created a singular profile containing a contrast matrix, which was used to calculate a singular image (SI) and singular value. To test the stability of the singular profile, permutation testing was performed using 500 samples; to estimate pattern-wise and voxel-wise standard error, bootstrap resampling was conducted using 100 samples.

In all voxel-wise analyses, maps were created expressing the ratio of voxel salience over the bootstrap standard error (i.e., BSR) to identify statistically reliable relationships between individual brain voxels and the singular profile. To characterize the spatial distribution of voxel responses, we inspected BSR maps for clusters of reliably differentiated voxels, defined as 12+ contiguous voxels of BSR 2.81 or higher with a minimum peak of BSR 3.29 (approximately corresponding to a minimum spatial extent of 324 mm^3^, a 99.9% peak confidence interval and a 99% extent confidence interval). To better characterize large clusters, we also recorded within-cluster local maxima. Cluster labels were obtained by transforming peak MNI coordinates into standardized Talairach coordinates using a best-fit icbm2tal transform (Lancaster et al., 2007) and localizing these in a Talairach brain atlas (Mai, Paxinos, & Voss, 2007).

We also performed a variation on this technique whereby non-rotated PLS was performed on each of the two component contrasts to obtain resampled SIs for each contrast (Poppenk, McIntosh, Craik, & Moscovitch, 2010). We first sampled groups of participants with replacement 100 times to generate distributions of SIs for the two contrasts. Then, we correlated each pair of SIs-employing the same bootstrapped sample to generate the two SIs within each pair-to generate a distribution of correlation coefficients describing the relationship of the two SIs. Finally, we evaluated the stability of the SI-pair correlations as the ratio of the mean correlation coefficient over the standard error from the correlation coefficient distribution (i.e., bootstrap ratio; BSR). To evaluate voxel-wise effects, we took the product of the 100 SI-pair product images, then took the square root of the product images to transform values back into linear space. We divided the mean square root product image by the standard error square root product image to obtain a map expressing BSR values, which we inspected using the same voxel-wise procedure described above.

## Results

We inspected behavioral responses from the initial exposure of repeated proverbs (Phase 1) to confirm engagement, but of focal interest in our analysis were neural responses to novel, repeated and previously known proverbs during the proverb viewing phase (Phase 2). In all of our analyses, *n*=17. Because we were unable to match the task or perceptual format of previously known proverbs to that of their earlier exposures, we also altered task and perceptual delivery across exposures of repeated proverbs to control these factors. As such, our procedure was optimal for RS analysis, but was not suitable for analysis of item-wise savings time.

### Behavioral results

#### Task engagement

For most proverbs in Phase 1, participants made button presses to signify proverb comprehension (*M*=0.85, *SD*=0.08), and guess each proverb’;s cultural origin (*M*=0.90, *SD*=0.08). Participants also responded to most stimuli in the three Phase 2 tasks (target age task *M*=0.97, *SD*=0.02; quality task *M*=0.97, *SD*=0.02; baseline task *M*=0.98, *SD*=0.03). Each task in Phase 2 was assigned different response keys, but perseveration errors were found in only five trials. Classifier analyses of fMRI data revealed statistically indistinguishable levels of Phase 2 task engagement across proverb types (as reported in Poppenk & Norman, 2012).

#### Prior knowledge

In a post-scanning knowledge questionnaire, participants confirmed advance knowledge of English but not Asian proverbs (rate of known proverbs:
English, *M*=0.90, *SD*=0.10; Asian, *M*=0.10, *SD*=0.10). As the questionnaire was self-paced, all items received responses. We excluded from each participant’s subsequent analyses all English and Asian proverbs that they reported as unknown and known, respectively.

#### Priming

We analyzed median RT of each condition in Phase 2. We observed shorter median RT for repeated than novel items overall (Fig. 2) in a bootstrapped repeated measures contrast, with a bootstrap ratio (BSR) of 7.44, P<0.001. We also observed shorter median reaction times for prior knowledge than novel items, BSR=14.28, *P*<0.001, and repeated items, BSR=3.17, *P*<0.005.

**Figure 2.**
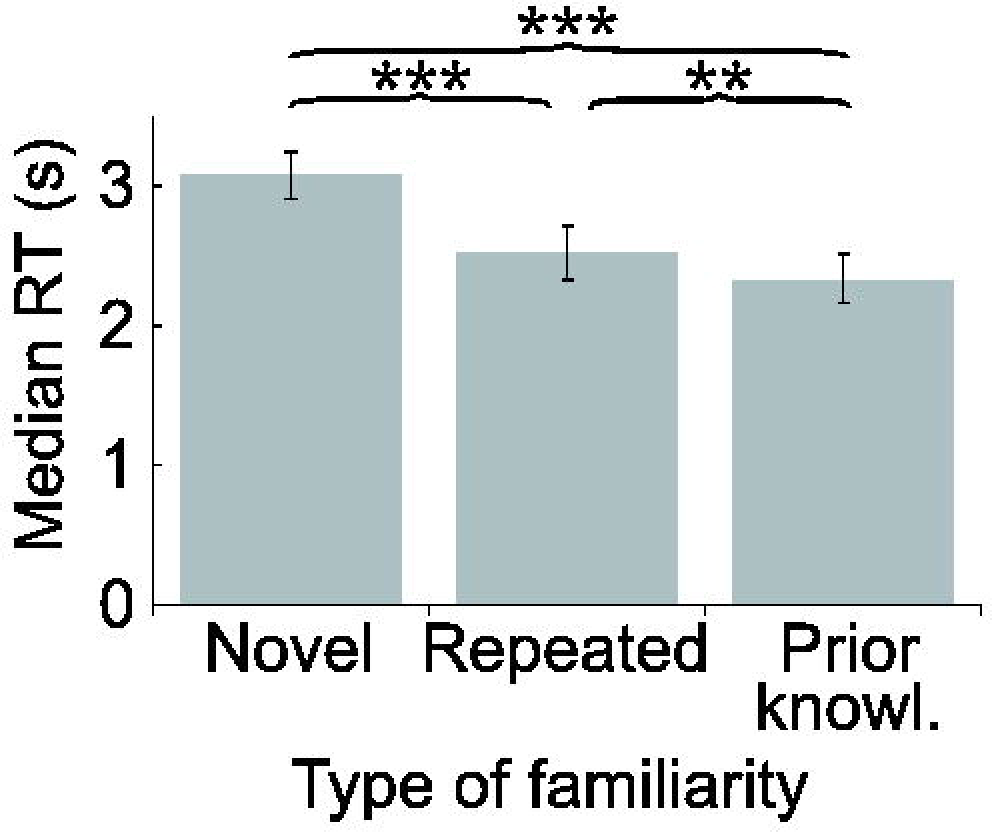
*LTM-based response facilitation*. Participants responded faster when experimental tasks involved proverbs in LTM (** *P* < 0.005; *** *P* < 0.001; error bars indicate 95% confidence intervals).

Savings time (i.e., a reduction in RT from Phase 1 to Phase 2) was not applicable to most proverbs, because only items from the repetition condition were presented by us in Phase 1 and therefore lacked any reference RT. Even in the case of repeated proverbs, the initial Phase 1 exposure involved a completely different task (is the proverb of African or East Asian origin?) than the one in Phase 2 (either the quality or target age task); therefore, even for this condition, there was no relevant Phase 1 reference RT. Consequently, we do not report savings time.

#### Source memory

Data from a source memory test that followed scanning are described elsewhere (Poppenk & Norman, 2012). Briefly, superior source memory was observed for prior knowledge proverbs relative to repeated ones, which in turn were associated with superior source memory relative to novel ones (see also Poppenk, Köhler & Moscovitch, 2010).

### Functional neuroimaging results

The objective of our fMRI analyses was to formally assess the possibility of: 1) a relationship in RS across LTM conditions; and 2) differences in RS strength across LTM conditions. Our principal tests were whole-brain multivariate analyses, but to better characterize relationships observed at the whole-brain level, we complemented these with descriptive voxel-wise analyses (see also McIntosh & Lobaugh, 2004). For reasons of statistical efficiency, we limited our exploration to one conjunction analysis evaluating the overlap of novelty vs. repetition and novelty vs. prior knowledge (i.e., testing RS stability across ~30-minutes and many-year repetition intervals); and one orthogonal comparison of repetition and prior knowledge (i.e., a test of RS decay and strengthening over the same period). We will henceforth use the term *LTM conditions* to describe these latter conditions, since both rely on forms of LTM.

#### Stable effects of repetition and prior knowledge

To determine whether each LTM condition engaged a common large-scale RS network, we conducted a multivariate conjunction test. In this test, we computed the correlation between the two whole-brain RS maps (repeated minus novel proverbs; and prior knowledge minus novel proverbs), taking the signal from all grey-matter voxels as input vectors. This test revealed a reliable pattern-level relationship between novel vs. repeated and novel vs. prior knowledge contrasts, *r*=0.65, BSR=8.32, P<0.001. We also performed a voxel-wise conjunction analysis using the same resampling procedure. All conjunction clusters were classified as either *stable RS* regions (repeated and prior knowledge < novel proverbs) or *stable RE* regions (repeated and prior knowledge > novel proverbs). We identified stable RS in visual and language cortices, including left iPFC and pSTG, and stable RE in frontal and parietal regions, including posterior cingulate and ventromedial prefrontal cortex (vmPFC; Figs. 3, 5; Table 3).

**Figure 3.**
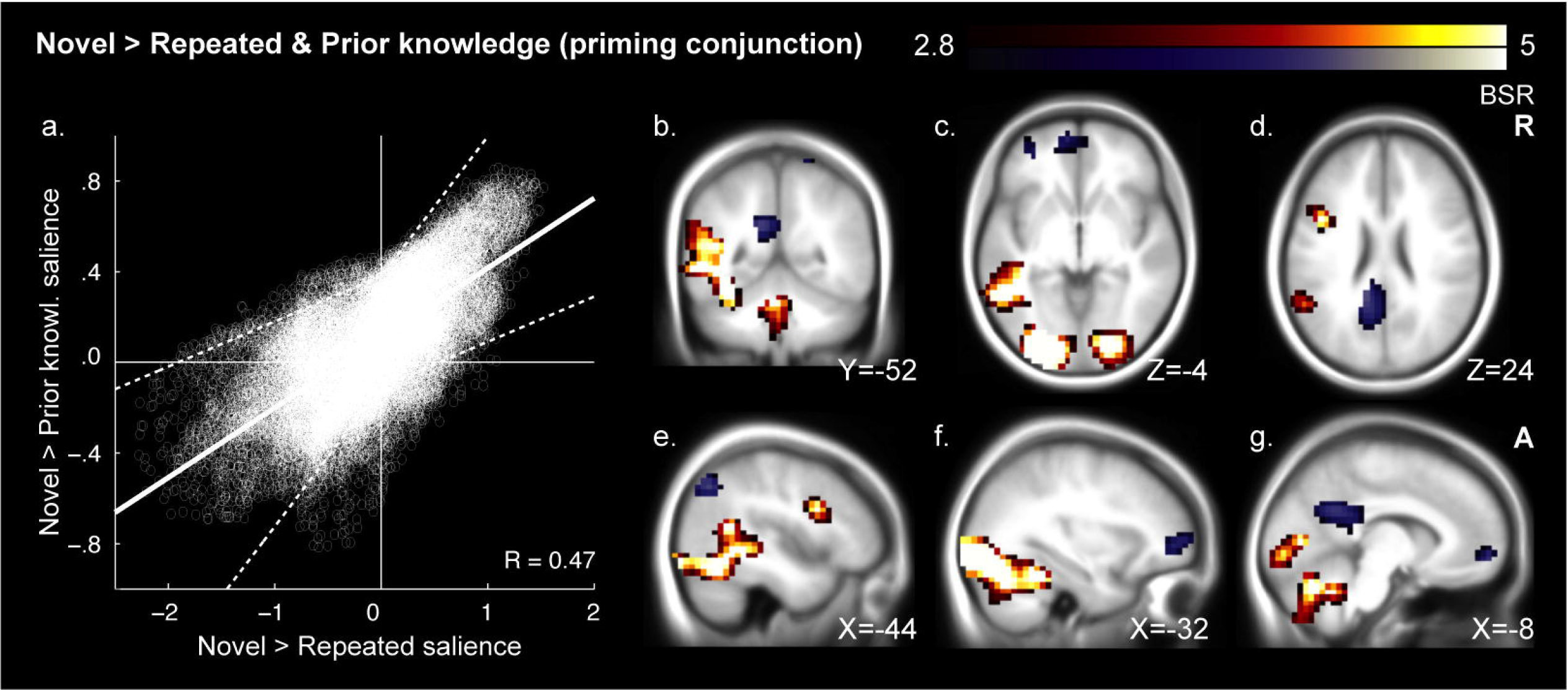
*Conjunction of repetition and knowledge-based RS*. Voxel saliences from the novel > repeated and novel > prior knowledge singular images were correlated (a). Their mean best linear fit following bootstrap resampling is shown as a solid line bounded by dashed 95% confidence limits. Their fit was visibly tighter in areas of RS (top/right) than stable RE (bottom/left). Reliable conjunction image voxels are presented on an average anatomical image of 200 young adults scanned at the Rotman Research Institute (b-g). Clusters contain peaks of at least BSR 3.29 and an extent of 324mm^3^ at a threshold of BSR 2.81 (corresponding approximately to 99.9% and 99.5% confidence thresholds). Orange depicts RS conjunction regions and blue stable RE regions. Priming conjunctions included perceptual-linguistic processing regions (b-g), whereas stable RE included areas frequently linked to memory retrieval (b-g). See Table 3 for regions and Fig. 5a for response plots.

**Table 3.** Voxel clusters associated with the conjunction contrast of novel vs. repeated and novel vs. prior knowledge proverbs.

| Region | BA | Hemi. | Peak MNI coordinates |  |  | BSR | Spatial extent (mm³) |
| --- | --- | --- | --- | --- | --- | --- | --- |
|  |  |  | X | Y | Z |  |  |
| Novel > Repetition & Prior knowledge ("Neural priming") |  |  |  |  |  |  |  |
| Frontal lobe |  |  |  |  |  |  |  |
| Broca's area | 44 | L | -44 | 8 | 24 | 5.07 | 3008 |
| Occipital lobe |  |  |  |  |  |  |  |
| Occipital g. | 17/18 | L | -20 | -88 | 4 | 8.64 | 127872 |
| Occipital g. | 17/18 | R | 20 | -88 | 0 | 8.06 |  |
| Posterior fusiform g. | 37 | L | -32 | -48 | -24 | 6.20 |  |
| Cerebellum | - | L | -16 | -72 | -36 | 10.22 |  |
| Cerebellum | - | R | 20 | -72 | -32 | 5.29 |  |
| Wernicke's area | 21/22 | L | -52 | -56 | 12 | 5.66 |  |
| Subcortical and other |  |  |  |  |  |  |  |
| Cerebellum | - | R | 20 | -36 | -36 | 3.49 | 576 |
| Repetition & Prior Knowledge > Novel ("Retrieval") |  |  |  |  |  |  |  |
| Frontal lobe |  |  |  |  |  |  |  |
| Medial frontal pole | 10 | L/R | -8 | 56 | -8 | 3.91 | 2048 |
| Lateral frontal pole | 10 | L | -32 | 48 | 0 | 3.74 | 2240 |
| Parietal lobe |  |  |  |  |  |  |  |
| Posterior cingulate g. | 23/31 | L/R | -12 | -44 | 24 | 4.15 | 7040 |
| Superior parietal l. | 7 | R | 20 | -52 | 72 | 3.30 | 832 |
| Angular g. | 39 | L | -44 | -68 | 40 | 4.07 | 3584 |

Note: Coordinates are displayed in MNI space.

#### Divergent effects of repetition and prior knowledge

It is important to note that independent of whether LTM conditions collectively differ from a novel baseline (as was the case among stable RS and RE regions), LTM conditions may also differ from one another, given that RS and RE may decay or strengthen between 30-minute and multi-year repetition intervals. This possibility was the basis of our second fMRI analysis goal, to probe for differences between the LTM conditions. A conventional non-rotated PLS analysis revealed that neural responses to repeated and prior knowledge proverbs could be readily distinguished at the whole-brain level, SV=5.44, P<0.005. Voxel-wise analysis revealed greater responses in vmPFC and left temporal pole to prior knowledge than repetition. In the repetition condition, we observed greater participation of lateral frontal pole, anterior and posterior cingulate cortex and a large set of parietal regions that included the precuneus and inferior parietal cortex (Figs. 4, 5; Table 4).

**Figure 4.**
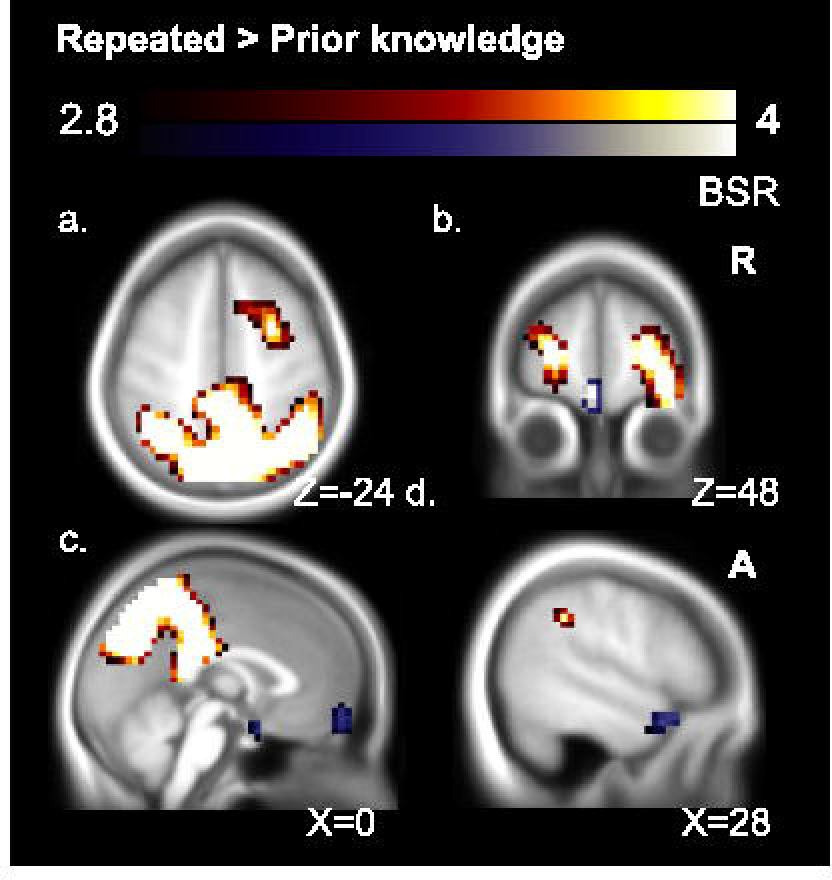
*RS differences*. Regions responding to repetition versus prior knowledge are overlaid on anatomy as in Fig. 3. Orange depicts repetition regions and blue prior knowledge regions. Repetition activations were observed in regions often found in episodic retrieval studies (a-c). Prior knowledge effects were seen in regions assocated with semantic memory, including ventomedial PFC (b,c), hypothalamus (c), and the left temporal pole (d). See Table 4 for regions and Fig. 5b for response plots.

**Figure 5.**
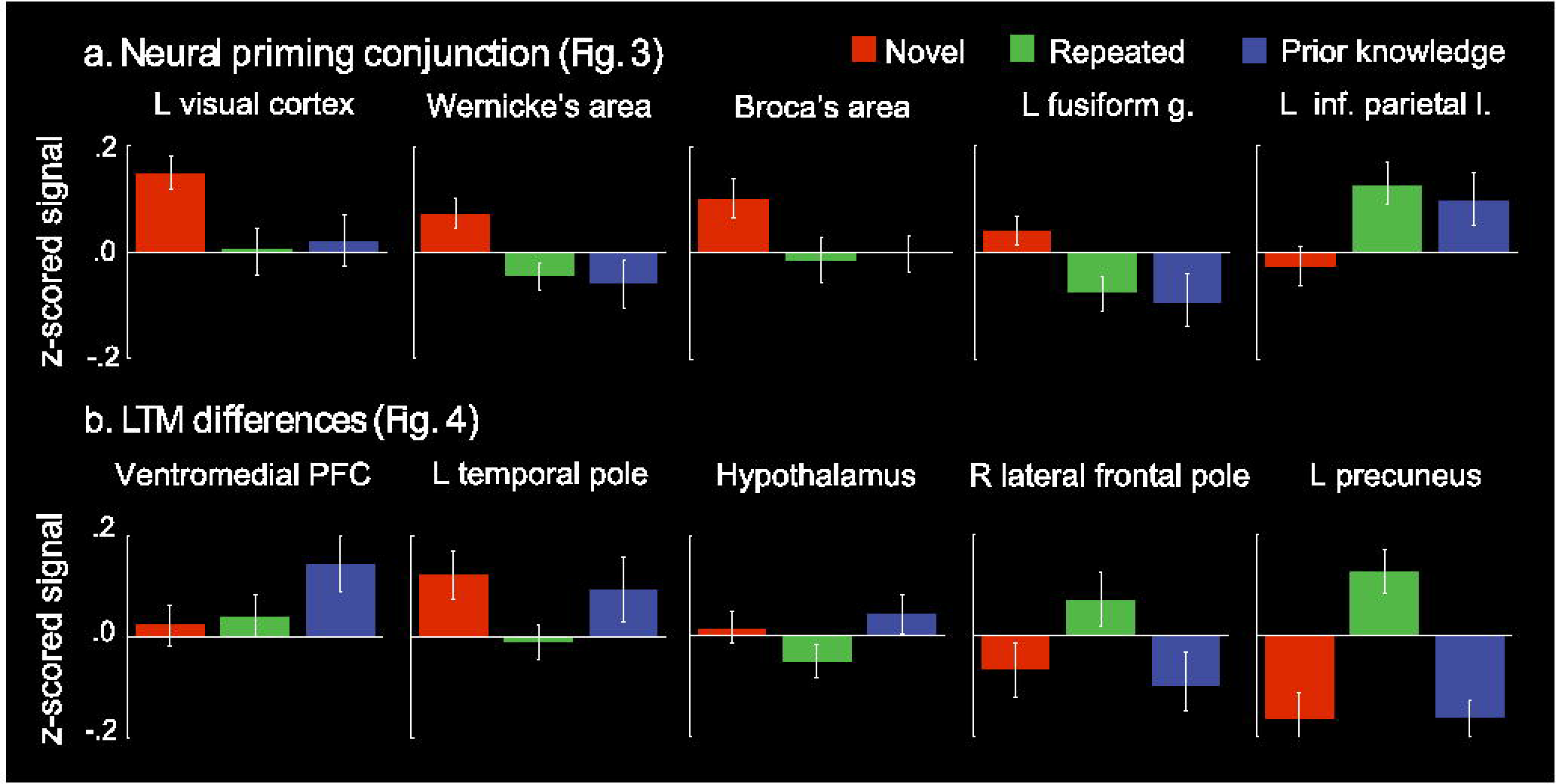
*Regional response plots for main contrasts*. Signal intensity in regions from the RS contrast (Fig. 3) is plotted with 95% confidence intervals about the mean (a). Similar plots are plotted for regions from the RS difference contrast (b; Fig. 4).

**Table 4.**
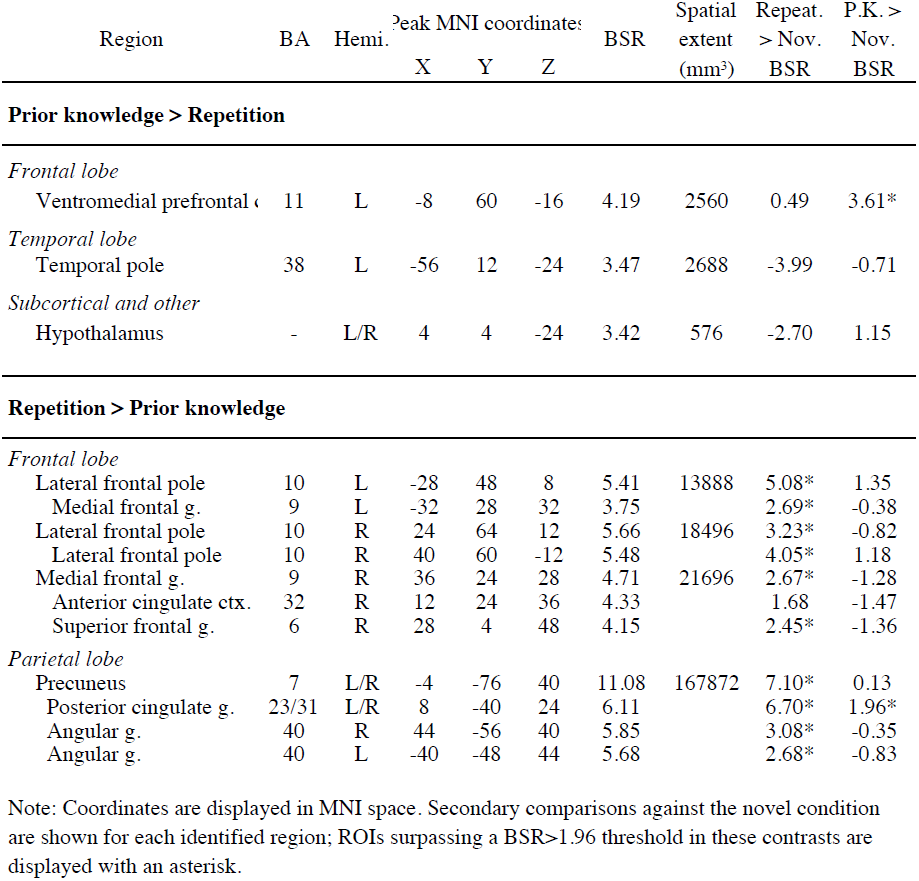
Voxel clusters associated with the contrast of repeated vs. prior knowledge proverbs.

Interestingly, while the above analysis showed that the LTM conditions differed, none of these differences fell within stable RS regions (i.e., that we identified in our conjunction analysis in Table 3). To confirm this observation, we conducted an ROI analysis, subjecting stable RS regions from the conjunction analysis (Table 3) to the contrast of repetition vs. prior knowledge using a threshold of BSR 1.96 (approximately corresponding to a 95% confidence interval). Even using this liberal test, we did not uncover any evidence of differences between the LTM conditions within RS regions. We did, however, identify differences between LTM conditions among three of the five stable RE regions (Table 3; Fig. 5).

The results described so far indicate that stable RS can be observed in broad portions of sensory and language cortices, with no difference in strength discernable on the basis of whether items are recently repeated or previously known. However, we may have overlooked other regions showing selective RS or RE to only one LTM condition. Because such regions are expected specifically in brain areas where LTM differences are found, we tested all of the fourteen LTM difference peaks from Table 4 for selectivity to one of the pairwise novelty contrasts, defined as supra-threshold (BSR=1.96) significance for one but not both of the repeated > novel and prior knowledge > novel contrasts. We found selective RS to repetition in the hypothalamus and left temporal pole (see Table 4, rightmost columns). We also observed selective RE to prior knowledge in vmPFC, consistent with studies linking vmPFC to schematic retrieval (van Kesteren, Rijpkema, Ruiter, & Fernandez, 2010). Among ROIs selected for greater repetition than prior knowledge activation, RE regions were consistently selective to repetition (Table 4, rightmost columns; with the borderline exception of posterior cingulate cortex), consistent with the possibility of episodic retrieval mechanisms evoked specifically by recently encoded proverbs.

#### Early and late repetition

As a follow-up analysis, we returned to whole-brain comparisons to test whether differences in RS could be observed within the repetition condition from the first half of the experiment (mean repetition interval = 20.4 minutes) to the second half of the experiment (mean repetition interval = 44.6 minutes; i.e., *RS decay*). Using non-rotated partial least squares, we tested an interaction crossing RS and time. This pattern-level effect was not significant, *P* = 0.81, suggesting that even at a repetition interval as short as 20 minutes, RS was relatively stable.

#### Large-scale shift in functional connectivity

We next explored whether RS in the two LTM conditions was linked to a shift in functional connectivity, as might be expected if local information processing were facilitated by information communicated by regions involved in LTM retrieval. To this end, we performed the same pattern-level conjunction and LTM difference analyses described above, but used seed-based covariance data in place of mean responses (Table 5). In particular, we used as seeds the peaks of each stable RS region reported in Table 3. In each region, we observed a common large-scale shift in functional connectivity in both LTM conditions relative to the novel condition, consistent with the possibility that RS in those regions arose on the basis of information communicated from elsewhere in the brain. Tests for large-scale RS differences in functional connectivity, which could indicate different overall patterns of regional communication for the two LTM conditions, were generally consistent, with only left iPFC, left pSTG and right cerebellum exhibiting trends towards such an effect (Table 5).

**Table 5.**
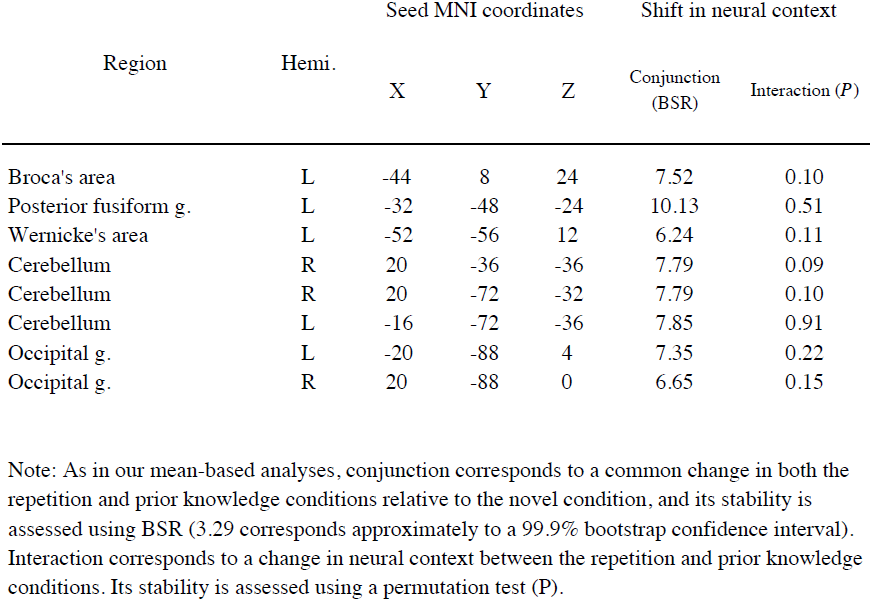
Retrieval-based changes in neural context associated with ROIs identified as neural priming regions (Table 3).

In the above functional connectivity analysis, it is worth noting that we selected seeds based on their association with the same hypothesized design we evaluated in covariance space, raising the risk that our result could have been pre-determined by our seed selection strategy. To test this possibility, we recorded the relationship between the covariance SIs for each pair of seeds. If our result was determined by the mean response that formed the basis of seed selection, spatial patterns associated with the seeds should share a large portion of their variance. However, the greatest R^2^ was 0.40 and mean R^2^ was 0.19 (Fig. S1 [suggested supplemental materials]), indicating our voxel selection strategy did not pre-determine the outcome of our seed covariance analyses.

#### Hippocampal region of interest

Returning to mean-based analysis, hippocampal suppression has been observed in response to stimulus repetition (e.g., Poppenk, McIntosh, et al., 2010; Tulving, Markowitsch, Craik, Habib, & Houle, 1996); we did not, however, identify any hippocampal region in our analyses. To test whether any effects were present but not detected under our statistical criteria, we searched for stable RS, stable RE and LTM difference voxel clusters within the boundaries of the hippocampus using a liberal threshold of BSR 1.96. While we still did not identify any stable RS region of the hippocampus, we did observe an LTM difference region: in particular, we observed greater activation of the left anterior hippocampus to prior knowledge proverbs than repeated ones, xyz=[−24 −16 −20], BSR=2.46, extent at BSR 1.96=2112mm^3^. This difference likely reflected RS that was selective to the repetition condition, as activity in this same ROI was suppressed in the repetition relative to novel condition, BSR=2.54, whereas no such difference was found between the prior knowledge and novel conditions, BSR=−0.43.

## Discussion

While RS effects fade rapidly at first, past evidence hints at RS stability at intervals lasting minutes to days. We assessed a high upper bound for such RS persistence by comparing suppression induced by recently repeated verbal materials and well-known, and long-ago repeated, and likely long dormant verbal materials that were matched in their properties. We found reliable large-scale overlap of RS, with voxel-wise analysis implicating a broad network of left-lateralized sensory and linguistic cortices (including occipital and fusiform gyri as well as Wernicke’;s and Broca’;s areas). No differences in RS strength were found among any of these common RS regions, and only two other regions (the hypothalamus and left temporal pole) showed RS selectively to one stimulus type (recent repetition). In addition, at each node of the shared RS network, presentation of either previously known or recently repeated proverbs induced the same shift away from the neural context associated with novel proverbs, further underscoring the similarity of the RS phenomenon induced by the two stimulus types. Because nearly identical neural suppression and connectivity changes were observed in response to highly remote and recently repeated stimuli, our evidence indicates that neural suppression characteristic of RS has an extremely long persistence.

In contrast to the striking homogeneity seen in most RS regions, almost all RE regions (e.g., vmPFC, temporal pole, and precuneus) were differentially associated with repetition and prior knowledge. In particular, prior knowledge was associated with selective RE in vmPFC, a region which may be crucial for retrieval of consolidated semantic, and possibly episodic, knowledge (van Kesteren et al., 2010); and the temporal pole, thought to constitute a hub that coordinates distributed elements of semantic information in cortex (Patterson, Nestor, & Rogers, 2007; see Liu, Grady and Mosovitch, in press). By comparison, repetition was associated with selective RE in “core” regions observed in episodic memory retrieval studies (Spaniol et al., 2009; Svoboda, McKinnon, & Levine, 2006), including precuneus, anterior and posterior cingulate gyrus, inferior parietal lobe, vmPFC and lateral PFC (but not the hippocampus). These differences in RE cannot be attributed to perceptual, task or motor priming effects specific to the repetition condition (Schacter et al., 2007), because the prestudy repetition and study phases featured different tasks, responses and fonts. Instead, because infrequent use of English proverbs means their presentation more likely triggers semantic than episodic memory, and because the opposite is true of repeated Asian proverbs, where insufficient time for consolidation has elapsed, this dissociation of RE regions most likely reflects the engagement of semantic and episodic memory networks for prior knowledge and recently repeated stimuli, respectively.

Hippocampal RS, especially in the anterior portion, has often been found in repetition studies similar to this one (Daselaar, Fleck, & Cabeza, 2006; Poppenk et al., 2008; Poppenk, McIntosh, et al., 2010; Strange & Dolan, 2001; Tulving et al., 1996). While we did not observe any hippocampal effects in our whole-brain contrasts, a liberal search for novelty-sensitive hippocampal voxels revealed a left anterior hippocampus region specifically suppressed by repetition, replicating earlier anterior hippocampal RS findings. This result extends earlier findings by showing that the anterior hippocampus, unlike most RS regions we observed, is responsive to cues related to recent episodes but not to those related to prior knowledge. This result is consistent with the structure’;s putative role in episodic, but not semantic, LTM (Moscovitch et al., 2005; Squire & Bayley, 2007). While the result is not surprising in light of past findings, it is interesting to juxtapose the involvement of the hippocampus in the set of regions suppressed by recent exposure, whereas the remainder of the episodic memory network was implicated in an RE network. While it is not possible to ascertain the meaning of the different response orientation of these elements of the episodic memory network, it is possible to speculate that whereas the complete episodic retrieval network is needed to reinstate memories, reduced hippocampal activity may reflect reduced encoding demands: most of the episode has already been stored, and only updated elements need to be recorded.

Broadly speaking, the other regions associated with stable RS were ones thought to be implicated in visual and linguistic processing. In addition to visual cortex, we observed stable RS in left iPFC and pSTG, regions known for their phonological/syntactic and semantic contributions to language (Rogalsky & Hickok, 2011; Vandenberghe, Nobre, & Price, 2002). We also observed stable RS in left posterior fusiform gyrus, a region believed to link meaning to word form or other visual information (Devlin, Jamison, Gonnerman, & Matthews, 2006), and in bilateral cerebellum, which also has been linked with linguistic function (Murdoch, 2010). No difference in RS strength could be detected in any of these regions. Previous repetition manipulations involving sentences have similarly evoked RS in the bilateral cerebellum, left iPFC, left pSTG, and primary sensory areas (Dehaene-Lambertz et al., 2006; Hasson et al., 2006). Our results show that such RS effects can be elicited not only by recent repetition, but also by prior knowledge.

Further to our finding of similar RS among recently repeated and remote stimuli, we observed no evidence of large-scale RS decay from the first to the second half of the experiment, similar to observations of little RS decay at one hour versus one day (van Turennout et al., 2003). These findings suggest that, in contrast to the period of rapid RS decay seen among repetition intervals of seconds to minutes (Henson et al., 2004; Henson et al., 2000; van Turennout et al., 2000; Wagner et al., 2000), RS for longer time intervals is relatively durable and stable. This pattern is consistent with the view that different RS mechanisms come into play after several minutes (Epstein, Parker, & Feiler, 2008; Weiner, Sayres, Vinberg, & Grill-Spector, 2010). One should be careful, however, not to interpolate between the time points represented by our prior knowledge and recent repetition conditions. The former likely reflects information consolidated into semantic memory from several exposures spaced over time, conditions known to lead to more robust representations (Spitzer, 1939). Based on the apparent importance of LTM in RS, it is possible, if not likely, that RS associated with materials that are not repeated or reinforced would decay at the same rate as corresponding episodic memories. In other words, the important contribution of our study was not to show that RS does not decay. Rather, it was to illustrate that typical RS can be evoked even under conditions where recent biological events or episodic memory are unlikely to play a role.

Although our study was a blocked design not suitable for direct correlation of neural RS and behavioral priming, facilitation of reaction times was observed behaviorally in both the prior knowledge and repetition conditions. Evidence from other studies also supports the idea that the phenomena are linked. For example, among studied items subsequently remembered on a recognition memory test, Turk-Browne and colleagues (2006) observed positive correlations between behavioral priming of scenes and response magnitude in regions implicated in processing them, such as bilateral PPA, fusiform gyrus (FFG) and right inferior PFC; however, these correlations were absent among subsequently forgotten items. Missing from the literature, however, has been any evidence that RS is as long-lasting as behavioral priming, which in previous (Kolers, 1976; Sloman et al., 1988), and more recent reports (Mitchell, 2006), has been shown to persist over long intervals of up to seventeen years. Because many years may go by between exposure to particular proverbs (see *Materials*), our results suggest that RS can have a persistence of similar magnitude. In this way, our results provide some support for the idea that RS and behavioral priming phenomena are connected.

These findings have implications for RS models. According to the sharpening account, cortical representations become sparser as information is consolidated into cortex (Wiggs & Martin, 1998). As systems consolidation is believed to operate over a period much longer than a single experimental session (Moscovitch et al., 2005), and as Phase II response times were shorter for prior knowledge proverbs than repeated ones, the prior knowledge proverbs in our study were likely consolidated better than the repeated ones. Accordingly, the sharpening model predicts greater sparsity of neural responses (and greater RS) for those proverbs. Nowhere in the brain was this pattern observed. In contrast, in the facilitation model, neural activity is reduced by top-down information from LTM, regardless of the age or source of memory. Our data fit this model better, as we observed comparable suppression of neural activity and shifting of functional connectivity in the two LTM conditions. That said, facilitation could still have led to reduced BOLD signal by way of either sparser neural responses or more rapid neural processing.

To reliably manipulate prior knowledge of materials in our experiment, our strategy was to juxtapose proverbs that are common in North America against ones more common in a different region. Although we matched these proverbs for length, and this approach allowed us to equate the literary quality of the materials, English and Asian proverbs may differ in other ways, raising the possibility that factors other than prior knowledge could have influenced our comparisons. Several pieces of evidence argue against this possibility. First, as illustrated in Fig. 3a, RS effects obtained by contrasting novel and familiar Asian proverbs were highly similar to those obtained in the contrast between novel Asian proverbs and prior knowledge proverbs. As Asian proverbs were randomly allocated to the novel and repeated conditions, it is unclear how cross-cultural effects could have led to conditions so similar to those of RS. Second, the brain regions identified in our contrast of repeated Asian and English proverbs were anticipated based on the predicted involvement of episodic and semantic memory in these conditions, respectively. These arguments notwithstanding, the current results, nevertheless should be considered preliminary, and call for further experimentation using other strategies to juxtapose RS resulting from recent stimulus exposure and RS resulting from prior knowledge.

In summary, we compared neural suppression induced by prior knowledge and recent repetition of verbal materials and observed suppression that was substantially equivalent in all forms in which we evaluated the effect. First, it was statistically indistinguishable in terms of magnitude and spatial extent within a large perceptual-linguistic network. Second, at each node of this network, previously known and recently repeated materials induced the same shift away from the neural context associated with novel materials. Accordingly, our results suggest RS is durable over intervals between about 30 minutes and many years. Also, remote and recently repeated proverbs most likely triggered retrieval from semantic and episodic memory, respectively, as ascertained by our finding of divergent RE activations that followed a spatial profile consistent with this dissociation. This finding, when coupled with stable RS, suggests that beyond merely being stable over time, RS is also stable across forms of LTM. Based on this pattern, as well as faster responding in both the prior knowledge and repetition condition, we propose that when repetition intervals exceed several minutes, RS is not a phenomenon related to decay that begins at the moment of initial stimulus exposure, such as one associated strictly with the episodic memory system or a separate short-lasting priming system. Rather, we propose that beyond initial, short period, RS reflects a general facilitation of perceptual and comprehension processes based on inputs from LTM, whether episodic or semantic, leading to faster and more accurate behavioral responses.

## Acknowledgements

We thank M. Ziegler for stimulus programming, F. Tam for imaging sequences, and N. Turk-Browne for helpful comments. Supported by NSERC-PGS (J.P.), NSERC-PDF (J.P.), NSERC-A8347 (M.M.), CIHR-MOP49566 (M.M), and J.S. McDonnell Foundation 22002082 (A.R.M.).

## Bibliography

Beckmann, C.F., & Smith, S.M. (2004). Probabilistic independent component analysis for functional magnetic resonance imaging. IEEE Trans Med Imaging, 23, 137–152.

Cocosco, C.A., Kollokian, V., Kwan, R.K. S., & Evans, A.C. (1997). Brainweb: Online interface to a 3D MRI simulated brain database. NeuroImage, 5, S425.

Craik, F. I. M., & Lockhart, R.S. (1972). Levels of processing: A framework for memory research. J Verbal Learn Verbal Beh, 11, 671–684.

Daselaar, S.M., Fleck, M.S., & Cabeza, R. (2006). Triple dissociation in the medial temporal lobes: recollection, familiarity, and novelty. J Neurophysiol, 96, 1902–11.

Dehaene-Lambertz, G., Dehaene, S., Anton, J.L., Campagne, A., Ciuciu, P., Dehaene, G.P., … Poline, J.B. (2006). Functional segregation of cortical language areas by sentence repetition. Hum Brain Mapp, 27, 360–71.

Devlin, J.T., Jamison, H.L., Gonnerman, L.M., & Matthews, P.M. (2006). The role of the posterior fusiform gyrus in reading. J Cogn Neurosci, 18, 911–22.

Eichenbaum, H., Yonelinas, A.P., & Ranganath, C. (2007). The medial temporal lobe and recognition memory. Annu Rev Neurosci, 30, 123–52.

Epstein, R.A., Parker, W.E., & Feiler, A.M. (2008). Two kinds of FMRI repetition suppression? Evidence for dissociable neural mechanisms. J Neurophysiol, 99, 2877–86.

Hasson, U., Nusbaum, H.C., & Small, S.L. (2006). Repetition suppression for spoken sentences and the effect of task demands. J Cogn Neurosci, 18, 2013–29.

Henson, R.N. (2003). Neuroimaging studies of priming. Prog Neurobiol, 70, 53–81.

Henson, R.N., Rylands, A., Ross, E., Vuilleumeir, P., & Rugg, M.D. (2004). The effect of repetition lag on electrophysiological and haemodynamic correlates of visual object priming. Neuroimage, 21, 1674–89.

Henson, R., Shallice, T., & Dolan, R. (2000). Neuroimaging evidence for dissociable forms of repetition priming. Science, 287, 1269–72.

Kolers, P.A. (1976). Reading a year later. Journal of Experimental Psychology: Human Learning and Memory, 2, 554.

Lancaster, J.L., Tordesillas-Gutierrez, D., Martinez, M., Salinas, F., Evans, A., Zilles, K., … Fox, P.T. (2007). Bias between MNI and Talairach coordinates analyzed using the ICBM-152 brain template. Hum Brain Mapp, 28, 1194–205.

Mai, J.K., Paxinos, G., & Voss, T. (2007). Atlas of the Human Brain (3rd ed.). San Diego, California: Academic Press.

McIntosh, A.R., & Lobaugh, N.J. (2004). Partial least squares analysis of neuroimaging data: applications and advances. NeuroImage, 23, S250–63.

Mitchell, D.B. (2006). Nonconscious priming after 17 years: invulnerable implicit memory? Psychol Sci, 17, 925–9.

Moscovitch, M. (1992). Memory and working-with-memory: A component process model based on modules and central systems. Journal of Cognitive Neuroscience. Special Issue: Memory Systems, 4, 257–267.

Moscovitch, M., Cabeza, R., Winocur, G., & Nadel, L. (2016). Episodic Memory and Beyond: The Hippocampus and Neocortex in Transformation. Annu Rev Psychol, 67, 105–34.

Moscovitch, M., Rosenbaum, R.S., Gilboa, A., Addis, D.R., Westmacott, R., Grady, C., … Nadel, L. (2005). Functional neuroanatomy of remote episodic, semantic and spatial memory: a unified account based on multiple trace theory. J Anat, 207, 35–66.

Murdoch, B.E. (2010). The cerebellum and language: historical perspective and review. Cortex, 46, 858–68.

Patterson, K., Nestor, P.J., & Rogers, T.T. (2007). Where do you know what you know? The representation of semantic knowledge in the human brain. Nat Rev Neurosci, 8, 976–87.

Poppenk, J., Kohler, S., & Moscovitch, M. (2010). Revisiting the novelty effect: When familiarity, not novelty, enhances memory. J Exp Psychol Learn Mem Cogn, 36, 1321–30.

Poppenk, J., McIntosh, A.R., Craik, F.I. M., & Moscovitch, M. (2010). Past experience modulates the neural mechanisms of episodic memory formation. J Neurosci, 30, 4707–16.

Poppenk, J., & Norman, K.A. (2012). Mechanisms supporting superior source memory for familiar items: A multi-voxel pattern analysis study. Neuropsychologia, 50, 3015–26.

Poppenk, J., Walia, G., McIntosh, A.R., Joanisse, M.F., Klein, D., & Kohler, S. (2008). Why is the meaning of a sentence better remembered than its form? An fMRI study on the role of novelty-encoding processes. Hippocampus, 18, 909–918.

Rogalsky, C., & Hickok, G. (2011). The role of Broca’;s area in sentence comprehension. J Cogn Neurosci, 23, 1664–80.

Schacter, D.L., Wig, G.S., & Stevens, W.D. (2007). Reductions in cortical activity during priming. Curr Opin Neurobiol, 17, 171–6.

Sloman, S.A., Hayman, C.A., Ohta, N., Law, J., & Tulving, E. (1988). Forgetting in primed fragment completion. Journal of Experimental Psychology: Learning, Memory, and Cognition, 14, 223.

Smith, S.M., Jenkinson, M., Woolrich, M.W., Beckmann, C.F., Behrens, T.E., Johansen-Berg, H., … Matthews, P.M. (2004). Advances in functional and structural MR image analysis and implementation as FSL. NeuroImage, 23 Suppl1, S208–219.

Spaniol, J., Davidson, P.S., Kim, A.S., Han, H., Moscovitch, M., & Grady, C.L. (2009). Event-related fMRI studies of episodic encoding and retrieval: meta-analyses using activation likelihood estimation. Neuropsychologia, 47, 1765–79.

Spitzer, H.F. (1939). Studies in retention. Journal of Educational Psychology, 30(9), 641–656.

Squire, L.R., & Bayley, P.J. (2007). The neuroscience of remote memory. Curr Opin Neurobiol, 17, 185–96.

Squire, L.R., & Zola, S.M. (1998). Episodic memory, semantic memory, and amnesia. Hippocampus, 8, 205–11.

Strange, B.A., & Dolan, R.J. (2001). Adaptive anterior hippocampal responses to oddball stimuli. Hippocampus, 11, 690–8.

Svoboda, E., McKinnon, M.C., & Levine, B. (2006). The functional neuroanatomy of autobiographical memory: a meta-analysis. Neuropsychologia, 44, 2189208.

Tulving, E., Markowitsch, H.J., Craik, F. I. M., Habib, R., & Houle, S. (1996). Novelty and familiarity activations in PET studies of memory encoding and retrieval. Cereb Cortex, 6, 71–9.

Turk-Browne, N. B., Yi, D.J., & Chun, M.M. (2006). Linking implicit and explicit memory: common encoding factors and shared representations. Neuron, 49, 917–27.

Vandenberghe, R., Nobre, A.C., & Price, C.J. (2002). The response of left temporal cortex to sentences. J Cog Neuro, 14, 550–60.

van Kesteren, M. T., Rijpkema, M., Ruiter, D.J., & Fernandez, G. (2010). Retrieval of associative information congruent with prior knowledge is related to increased medial prefrontal activity and connectivity. J Neurosci, 30, 15888–94.

van Turennout, M., Bielamowicz, L., & Martin, A. (2003). Modulation of neural activity during object naming: effects of time and practice. Cereb Cortex, 13, 381–91.

van Turennout, M., Ellmore, T., & Martin, A. (2000). Long-lasting cortical plasticity in the object naming system. Nat Neurosci, 3, 1329–34.

Wagner, A.D., Maril, A., & Schacter, D.L. (2000). Interactions between forms of memory: when priming hinders new episodic learning. Journal of Cognitive Neuroscience, 12, 52–60.

Weiner, K.S., Sayres, R., Vinberg, J., & Grill-Spector, K. (2010). fMRI-adaptation and category selectivity in human ventral temporal cortex: regional differences across time scales. J Neurophysiol, 103, 3349–65.

Wiggs, C.L., & Martin, A. (1998). Properties and mechanisms of perceptual priming. Curr Opin Neurobiol, 8, 227–33.

